## Supplementary material for "Super-silencers are crucial for development and carcinogenesis in B cells": supplmentary notes

### Supplementary Notes

#### Data for predicting silencers and identifying SSs.

We obtained TF ChIP-seq peaks for the GM12878 lymphoblastoid cell line as of April 2021 from the Encyclopedia of DNA elements project (https://www.encodeproject.org/) and DNase-seq peaks from the Roadmap Epigenomics Project (<http://egg2.wustl.edu/roadmap>/data/ byFileType/peaks/consolidated/broadPeak/). These peaks were combined as active genomic regions. To identify silencers, we screened these active genomic regions using a 1kb window and a step size of 100 bp with the CNN enhancer-silencer prediction model developed in our previous study ^1^.

To identify super-silencers (and super-enhancers), we downloaded GM12878 H3K27me3 and H3K27ac ChIP-seq alignment files from the Roadmap Epigenomics Project (<http://egg2.wustl.edu/roadmap>/data/byFile/Type/alignments/consolidated). Using the H3K27me3 ChIP-seq alignment file, we applied the ROSE program ^2^ to the predicted silencers, identifying the super-silencer (SS) regions and the silencers located within those regions (referred to as SS components). Similarly, the ROSE program was used with the H3K27ac ChIP-seq alignment file to detect the SEs and SE components among the predicted enhancers.

#### Data for evaluating the function of GM12878 silencers and enhancers across cell/tissue types.

We obtained H3K27ac and H3K27me3 ChIP-seq peaks from the Roadmap Epigenomics Project (http://egg2.wustl.edu/roadmap) for a total of 56 cell/tissue types, excluding GM12878 cell line. To characterize the function of the predicted GM12878 silencers/enhancers across these cell/tissue types, we overlapped the predicted silencers/enhancers with these H3K27ac and H3K27me3 ChIP-seq peaks. A silencer/enhancer was considered carrying H3K27ac/H3K27me3 modification (indicating the positive/negative regulatory function) when its midpoint fell within an H3K27ac/H3K27me3 peak.

#### Gene deserts.

Gene deserts were defined as over-500mb-long genomic regions that are absent of protein-coding genes^3^. Gene annotations used here were downloaded from the GENCODE project (gencode.v42.annotation.gff3.gz) ^4^.

#### Tissue-specificity of genes based on gene expression profiles.

We downloaded the RNA-seq data of 251 biosamples from the ENCODE project. The gene expression, measured as the Reads Per Kilobase of the transcript, per Million mapped reads (RPKM values), were normalized so that the expression level of each gene had a median of zero. Tissue-specificity of a gene was measured using tau (τ) ^5^ as

$\tau=\frac{\sum_{i=1}^{N} (1-\hat{x}_{i})}{N}, \hat{x}_{i}=\frac{x_{i}}{{max}_{k=1}^{N}x_{k}}$,

where $x_{i}$ is the expression of a tested gene in the cell line $i$. $N$ is the number of the cell lines under consideration. The values of $\tau$ are in the range of $[0,1]$. A high value of $\tau$ corresponds to a large variation in gene expression across tissues, i.e., a high tissue specificity. The genes with $\tau>0.97$ were considered as tissue-specific, while the genes with $\tau<0.8$ were labelled as housekeeping.

#### Silencers and enhancers in primary B cells and B-cell-cancer cell lines.

We obtained H3K27ac and H3K27me3 ChIP-seq peaks of primary B cells from the ENCODE project (biosample ID CL:0000788, https://www.encodeproject.org/biosamples/ ENCBS857XIR/). We applied our framework to these peaks, identifying silencers and SSs among them using the ROSE ^2^. The predicted silencers (TSs and SS components) are listed in Table S5.

In addition, we applied our framework to ChIP-seq data in six B-cell-cancer cell lines available in the ENCODE project, identifying silencers (and SSs) for these cell lines. These cell lines include Ly1 (ENCODE biosample ID EFO:0005907), Ly3 (EFO:0006710), Karpas-422 (EFO:0005719), and SU-DHL-6 (EFO:0002357, Table S4). Comparing silencers profiles in these cell lines with the enhancer profile in GM12878, we identified the conversions from SEs to SSs during the development of DLBCL.

### Supplementary Figures

#### Supplementary Figure 1

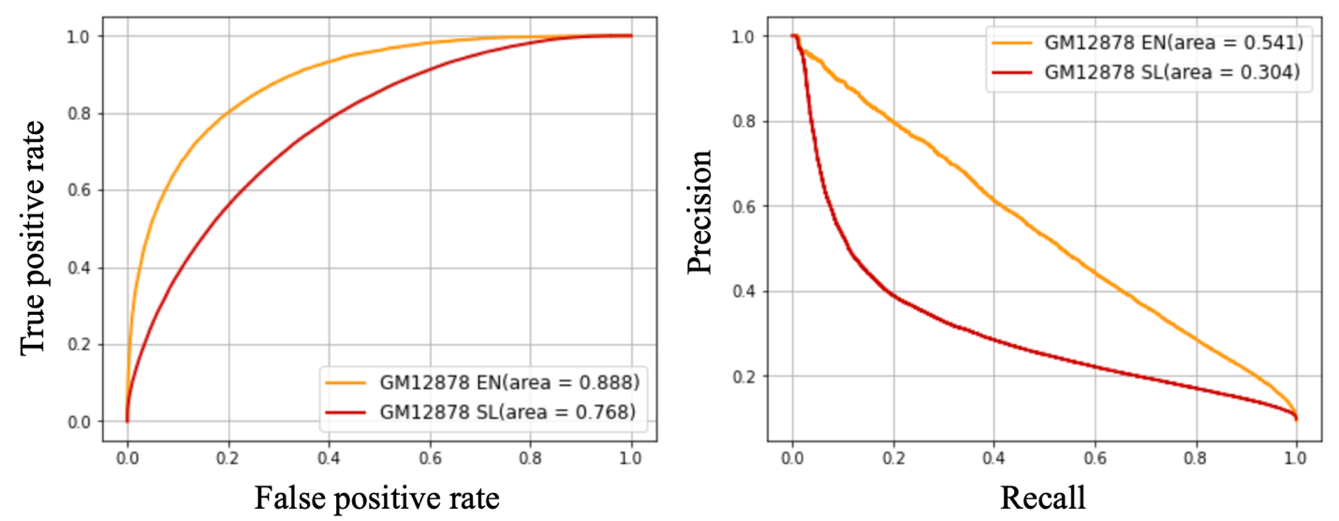

**Figure S1.** Performance of silencer and enhancer prediction in GM12878. The classifier here was built in our previous study ^1^.

#### Supplementary Figure 2

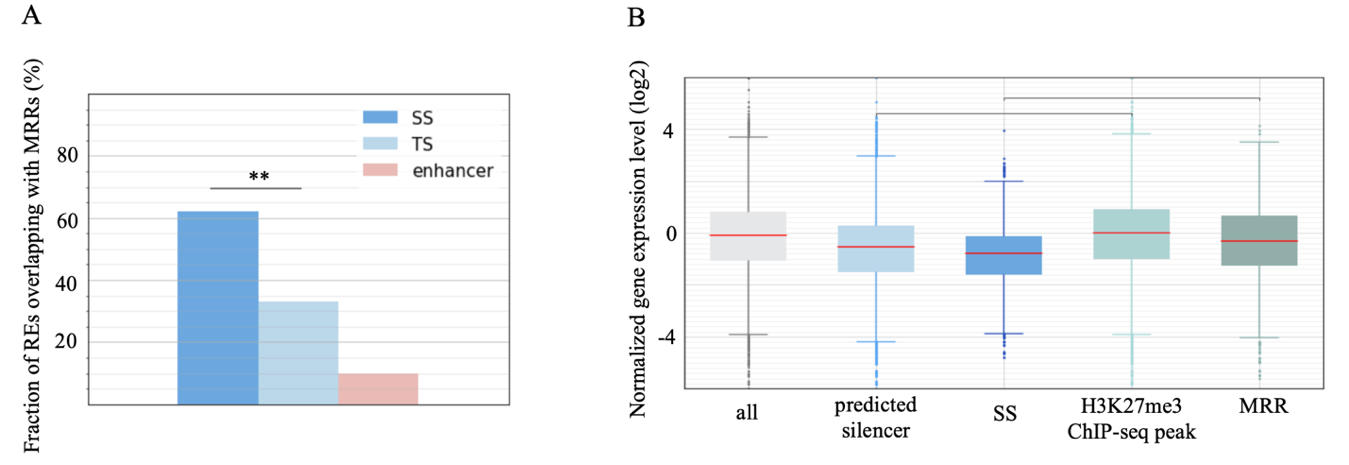

**Figure S2.** Overlap of the predicted SSs with MRRs in GM12878 ^6^. (A) Fractions of SS components, TSs, and enhancers overlapping with MRRs in GM12878. (B) Expressions of genes proximal to predicted silencers and SSs and MRRs.

#### Supplementary Figure 3

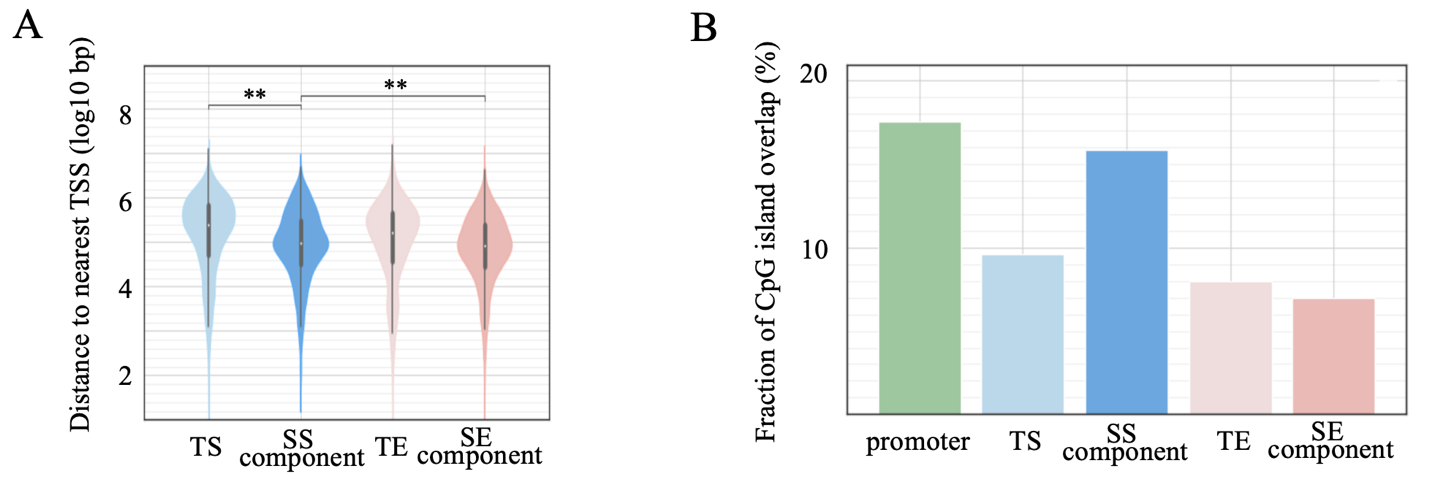

**Figure S3.** Genomic distribution of enhancers and silencers in terms of (A) distances to their most proximal TSSs and (B) overlaps of silencers and enhancers with CpG islands. $**- p<{10}^{-10}$.

#### Supplementary Figure 4

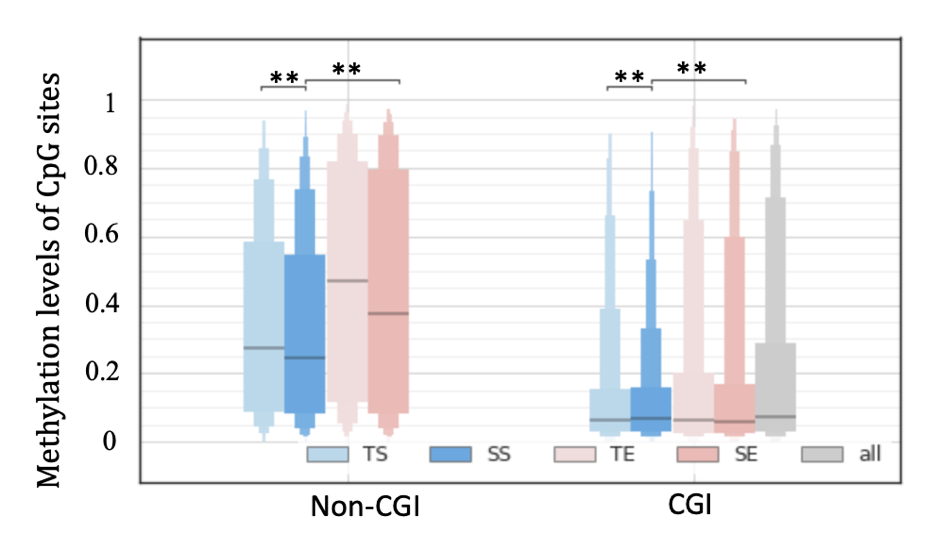

**Figure S4.** Methylation levels of CpG sites in enhancers and silencers. For a fair comparison, we subcategorized silencers/enhancers into CGI silencers/enhancers (i.e., silencers/enhancers overlapping CpG islands by >200 bp) and non-CGI silencers/enhancers.

#### Supplementary Figure 5

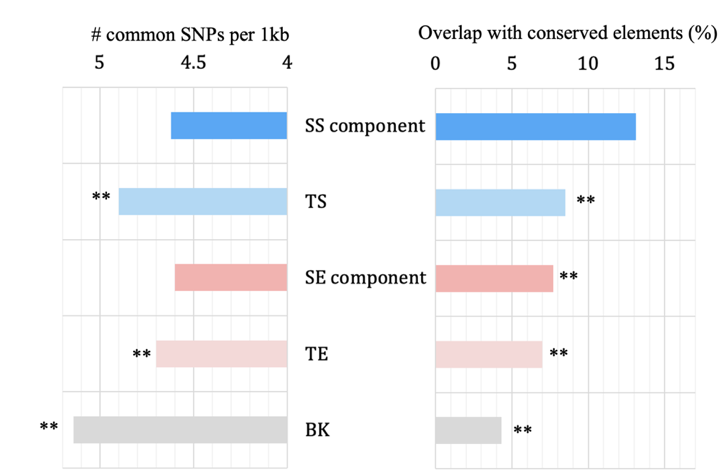

**Figure S5.** Conservation of enhancer and silencer types. (A) Numbers of common SNPs per 1kb. (B) Overlap with conversed elements. Conserved elements are detected based on Phastcon scores ^7^.

#### Supplementary Figure 6

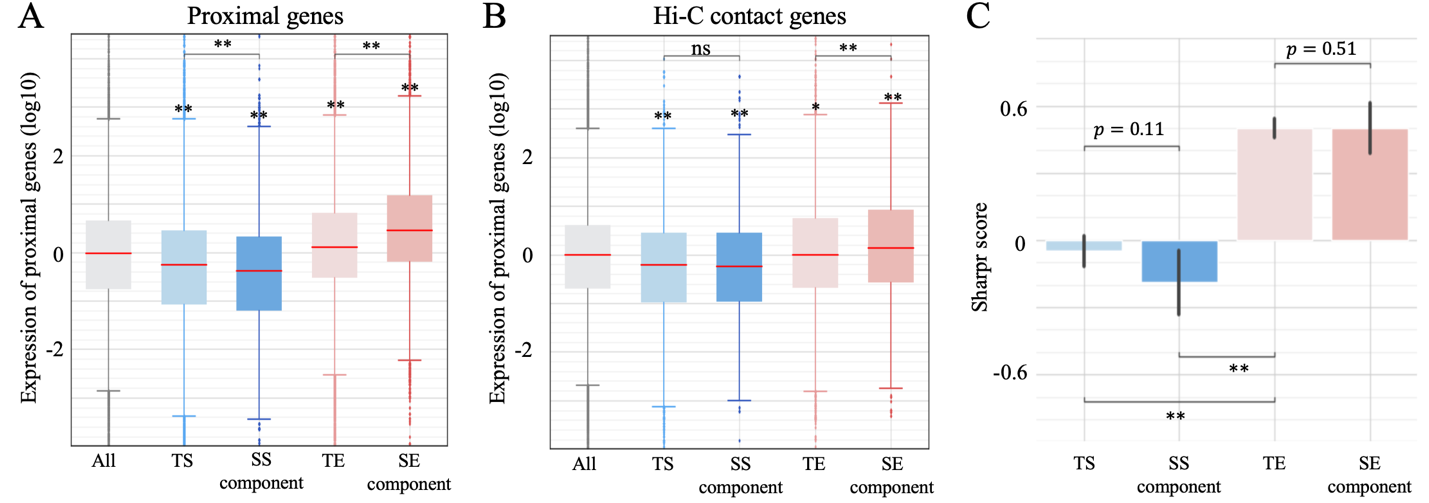

**Figure S6.** Evaluation of predicted super-silencers across different cell types, including hESC H1, HepG2 and K562. Our evaluation is based on expression levels of genes (A) proximal to regulatory elements and (B) having Hi-C contacts to regulatory elements, as well as (C) Sharpr scores of regulatory elements in HepG2 and K562 cell types.

#### Supplementary Figure 7

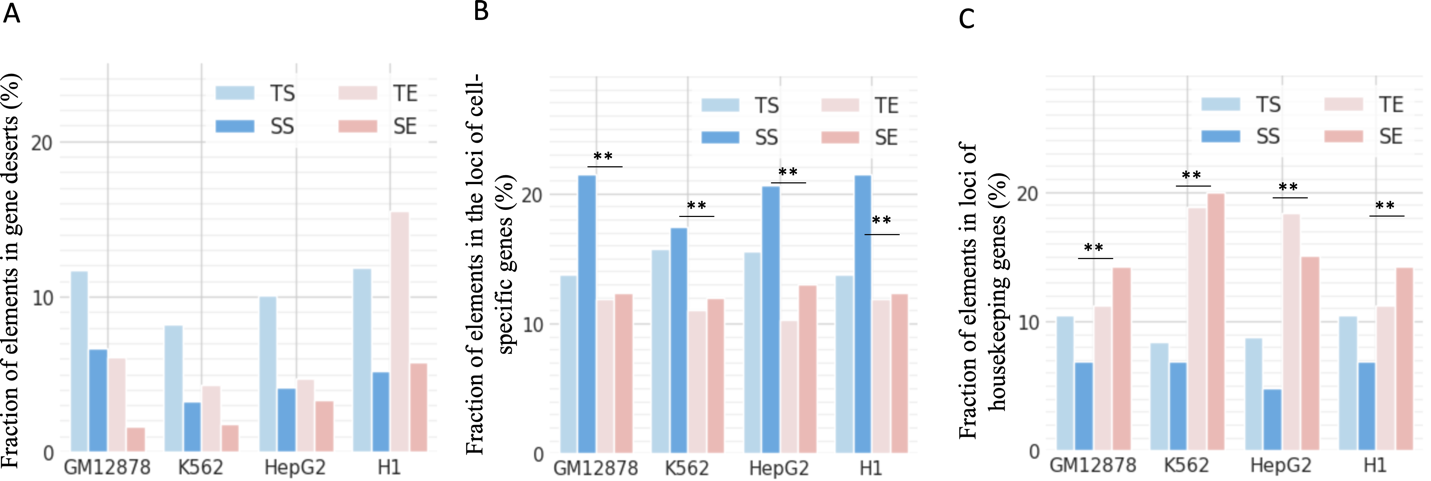

**Figure S7.** Distribution of predicted silencers and enhancers in the whole genome across different cell types. (A) Fraction of elements in gene deserts. (B) Fraction of elements of predicted super-silencers across different cell types, including hESC H1, HepG2 and K562. Our evaluation is based on expression levels of genes (A) proximal to regulatory elements and (B) having Hi-C contacts to regulatory elements, as well as (C) Sharpr scores of regulatory elements in HepG2 and K562 cell types.

#### Supplementary Figure 8

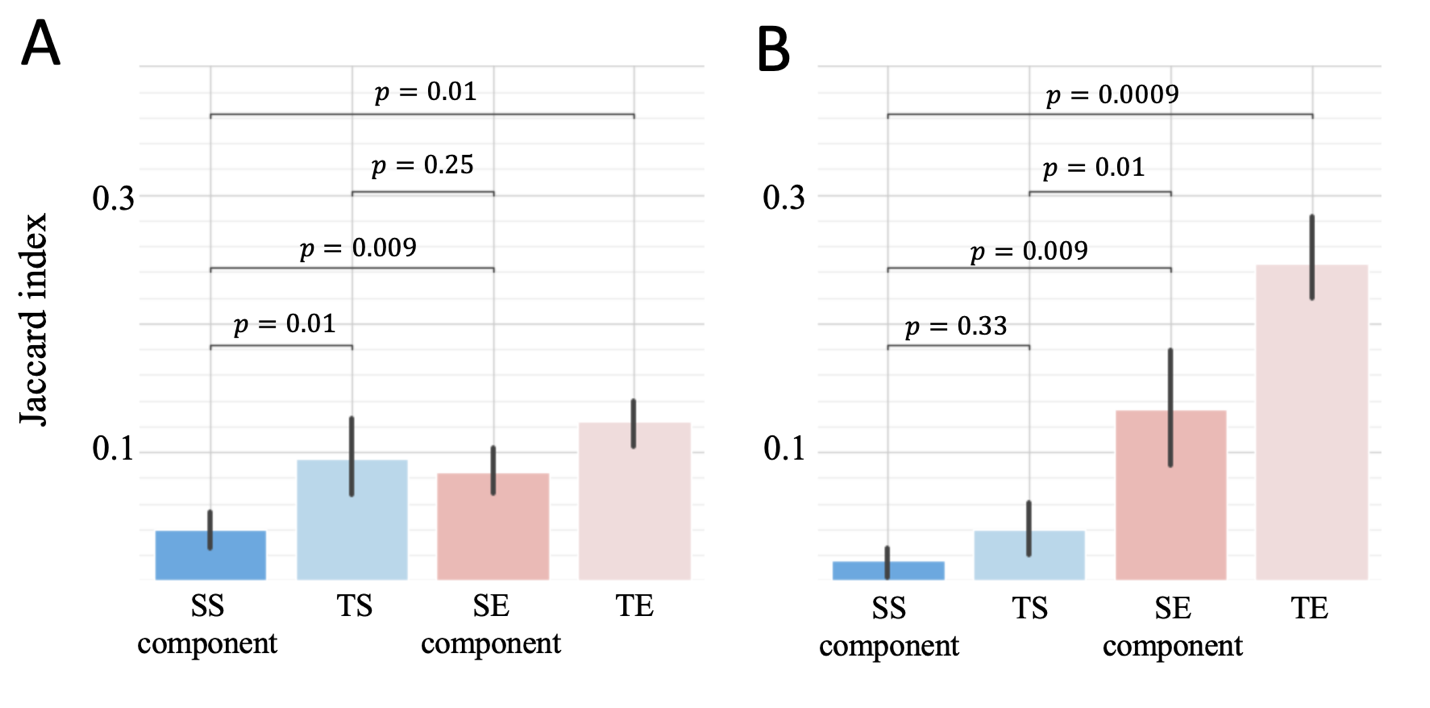

**Figure S8.** Similarities of regulatory element profiles across cell types. The similarity of silencer or enhancer profiles between two cell types was measured in terms of Jaccard index. (A) Similarities across four distinct cell types GM12878, K562, hESC H1 and HepG2. (B) Similarities between GM12878 and B cell lymphoma cell lines. Here, B-cell cancer cell lines include Ly1 (ENCODE biosample EFO:0005907), Ly3 (EFO:0006710), Karpas-422 (EFO:0005719), and SU-DHL-6 (EFO:0002357).

#### Supplementary Figure 9

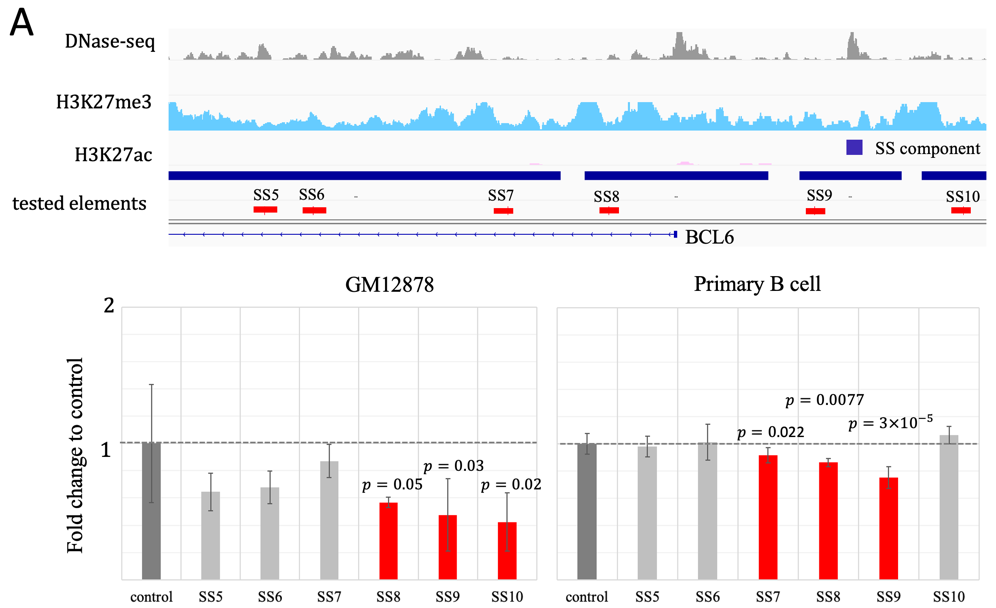

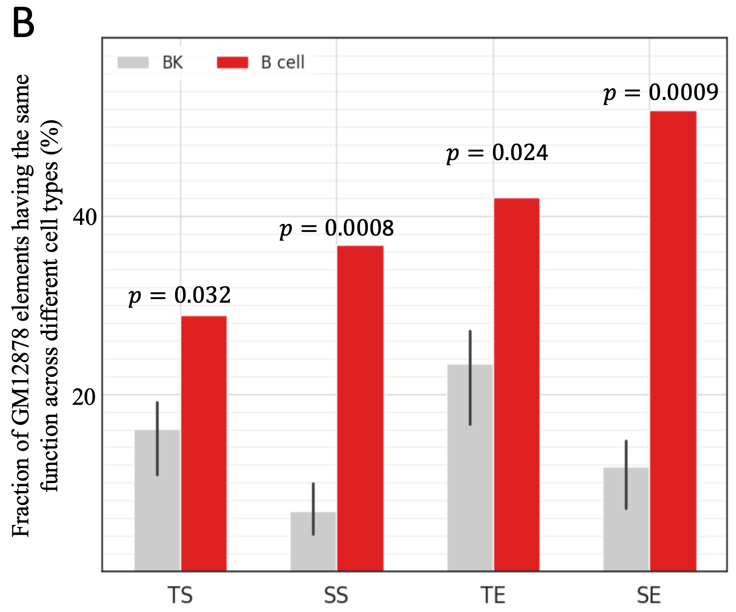

**Figure S9.** Experimental results of SS segments within the *BCL6* locus (A) in GM12878 and primary B cells. (B) Fractions of GM12878 silencers and enhancers displaying the same activity in other cell types. “BK” represents the cell types hESC H1, HepG2 and K562, while “B cell” represent the primary naïve B cells.

#### Supplementary Figure 10

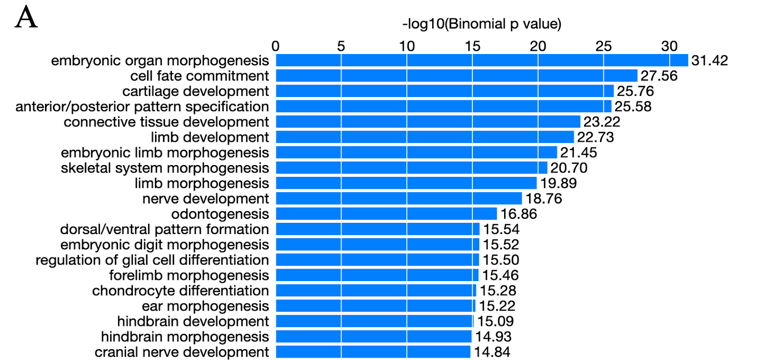

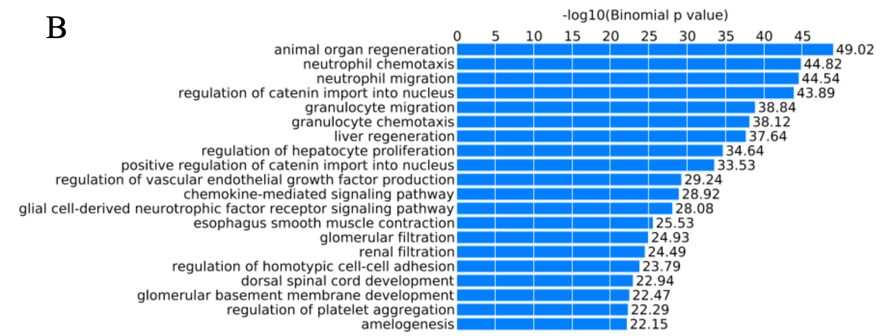

**Figure S10.** Function analysis of (A) GM12878 SS components and (B) TSs by using the webtool GREAT ^8^.

#### Supplementary Figure 11

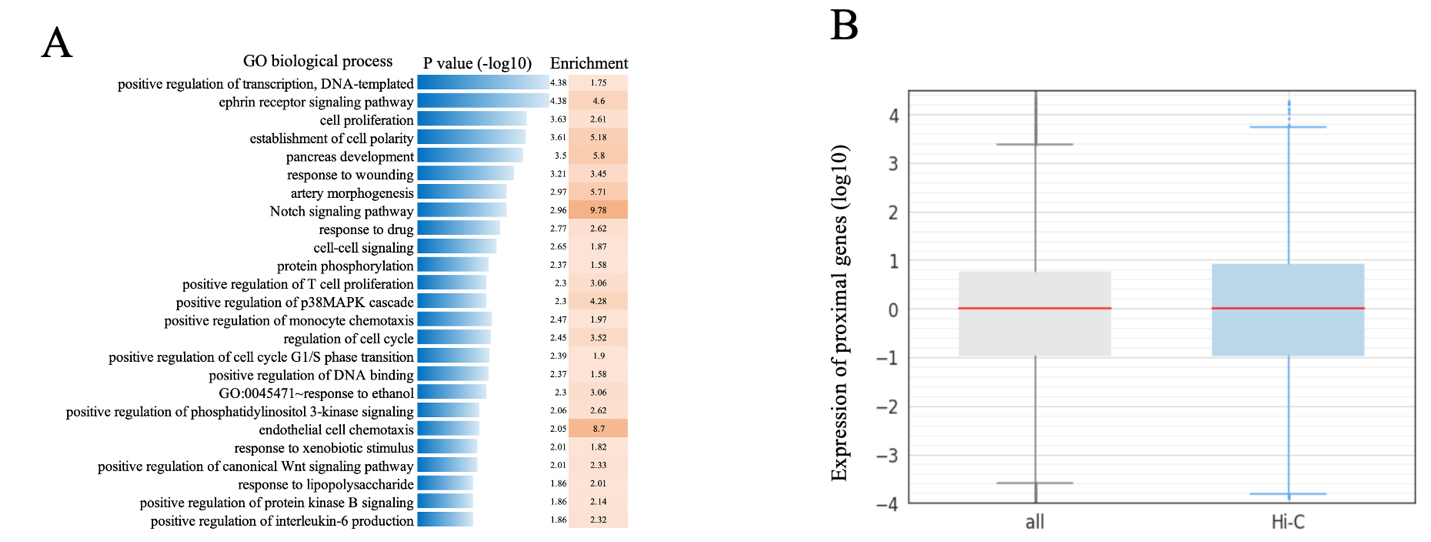

**Figure S11.** (A) Function analysis of genes interacting with GM12878 TSs by using the webtool DAVID ^9^. (B) Genes having Hi-C contacts show elevated expression levels compared to all genes.

#### Supplementary Figure 12

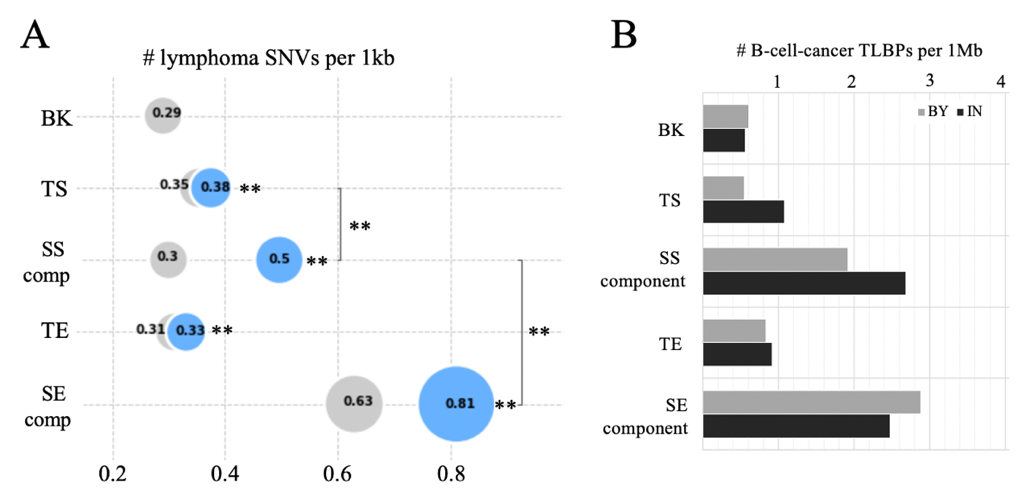

**Figure S12.** Distribution of cancer mutations in SSs. (A) Numbers of B-cell-cancer SNVs per 1kbp and (B) Numbers of B-cell-cancer TLBP per 1Mbp in silencers and enhancers.

#### Supplementary Figure 13

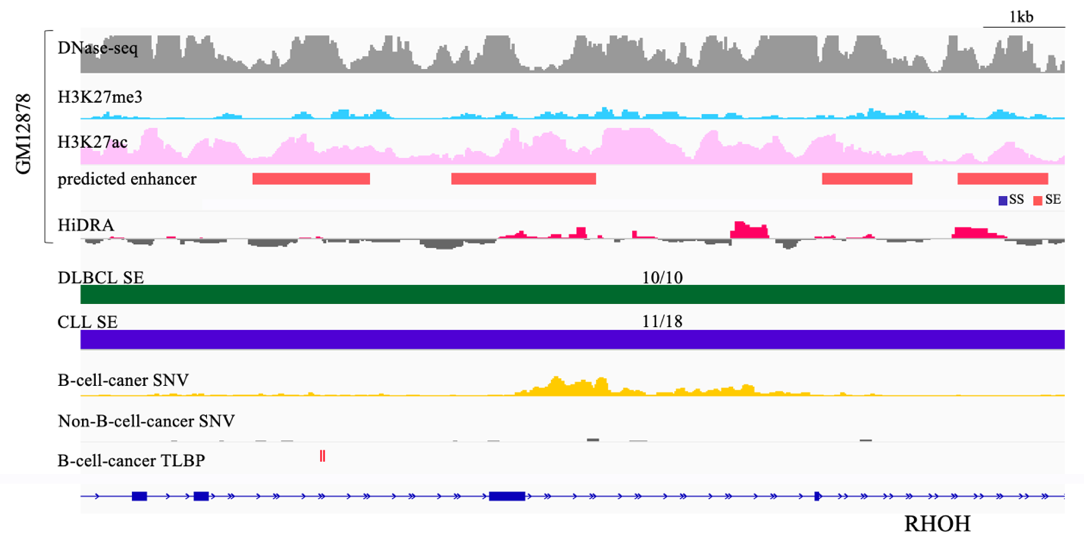

**Figure S13.** Genomic and epigenetic profiles of RHOH SSs in normal and cancer genomes.

#### Supplementary Figure 14

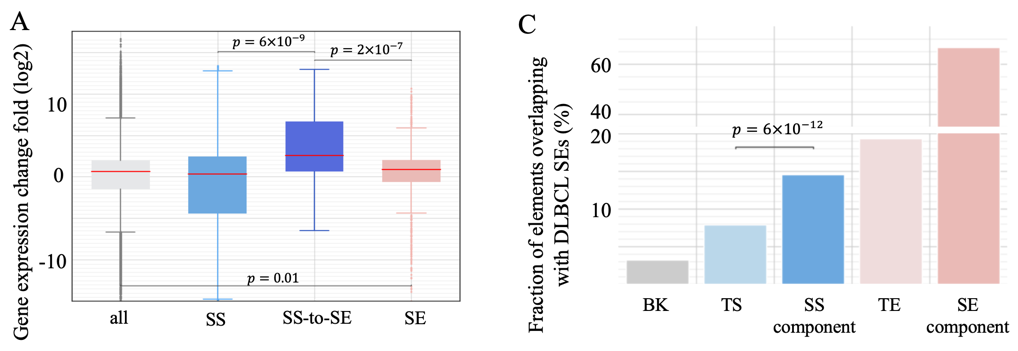

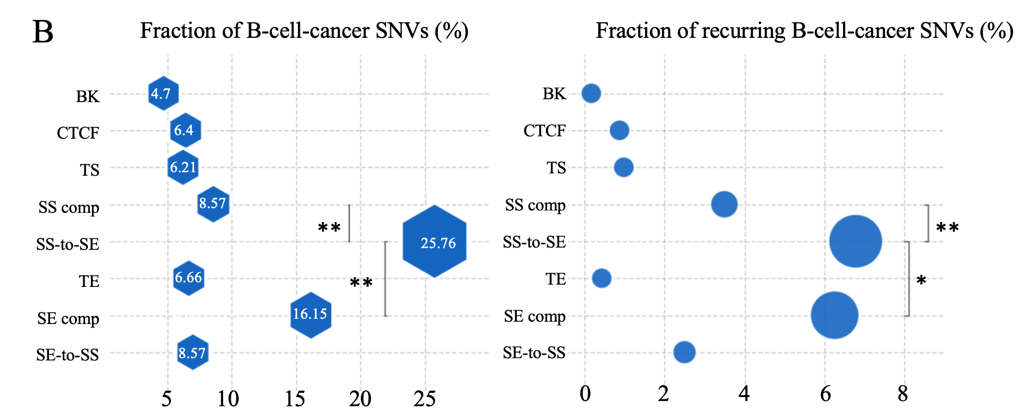

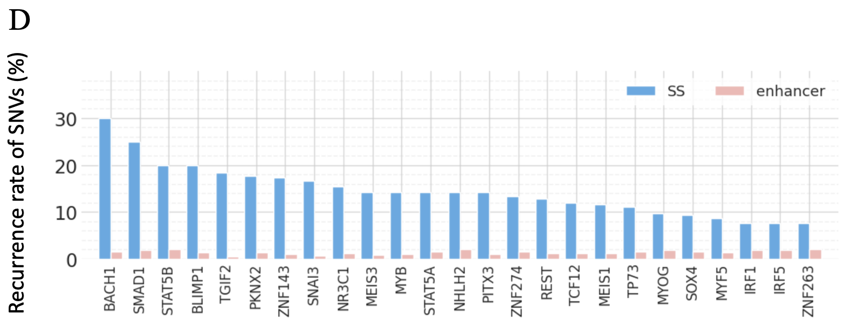

**Figure S14.** Impact of the conversion of SSs to SEs during DLBCL carcinogenesis. (A) Genes proximal to SS-to-SEs show the highest upregulation levels in DLBCL patients. (B) Enrichment of B-cell-cancer SNVs in SS-to-SEs. (C) High overlaps of SSs with DLBCL SEs reported by Elodie Bal et al. ^10^. (D) Fractions of recurring B-cell-cancer SNVs within binding motifs in SS-to-SE sequences.

#### Supplementary Figure 15

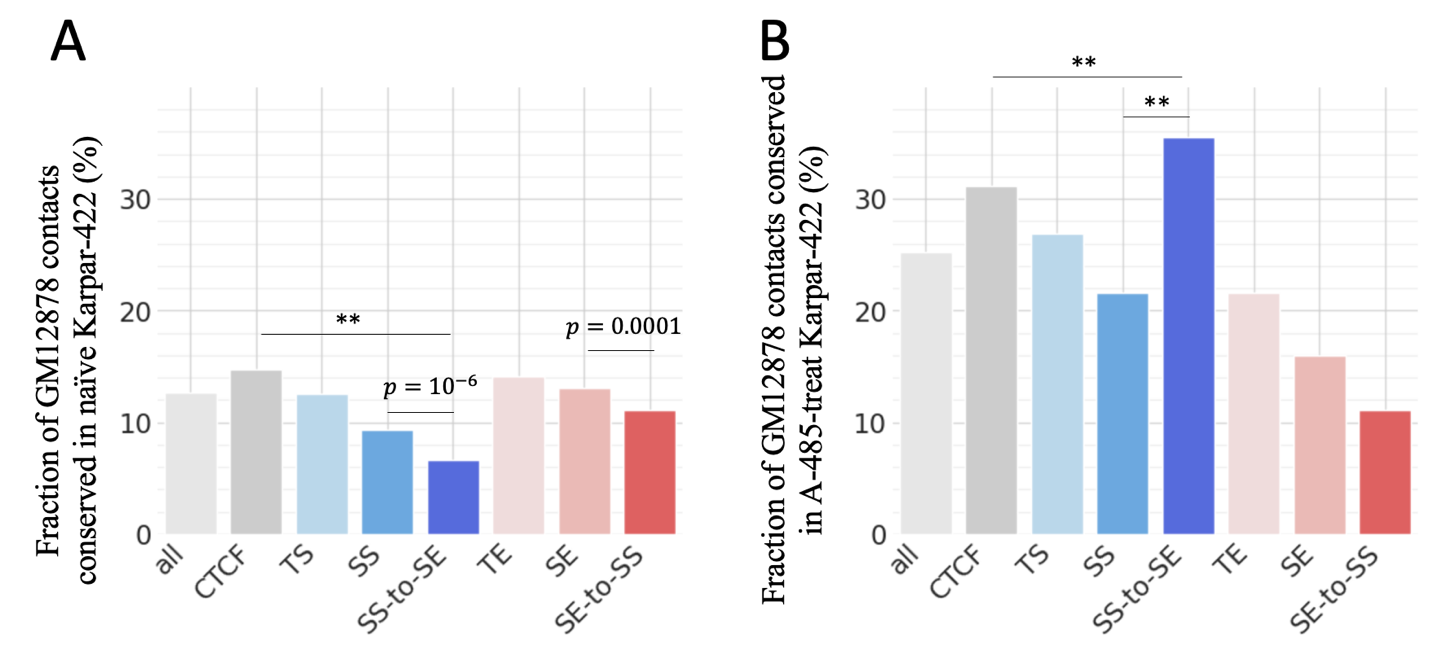

**Figure S15.** Chromatin structure changes in the B-cell lymphoma carcinogenesis. (A) Fractions of chromatin contacts conserved between GM12878 and Karpar-422, a B-cell lymphoma cell lines. (B) Overlap of GM12878 chromatin contacts with those in A-485-treat Karpar-422 cells. These overlap levels suggest the likelihood of chromatin contacts in normal B cell being restored by the A-485 treatment.

#### Supplementary Figure 16

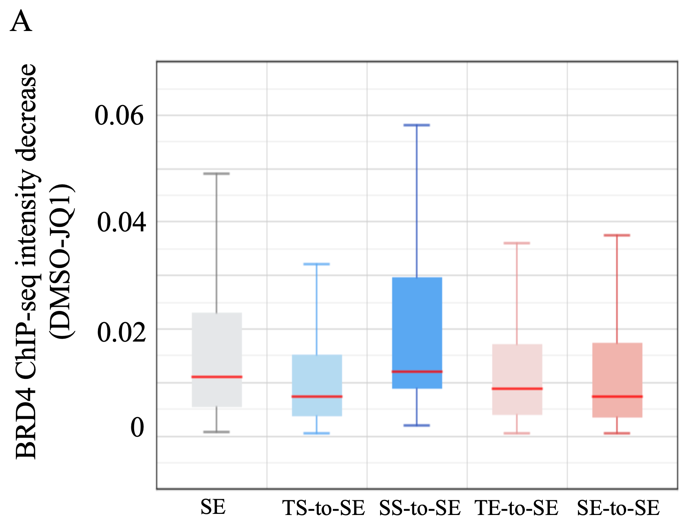

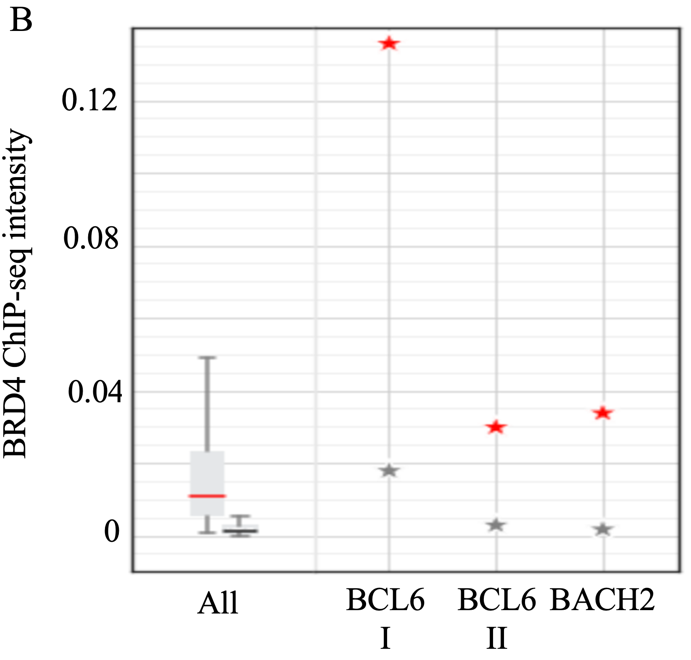

**Figure S16.** Change of BRD4 binding intensities (as measured by ChIP-seq signals) in DLBCL SEs after JQ1 treatment. (A) SS-to-SEs have the largest decrease in BRD4 binding intensities after JQ1 treatment. (B) BRD4 ChIP-seq signals in the DLBCL SEs within the *BCL6* and *BACH2*. Red and grey markers represent the signals before and after the JQ1 treatment, respectively.

#### Supplementary Figure 17

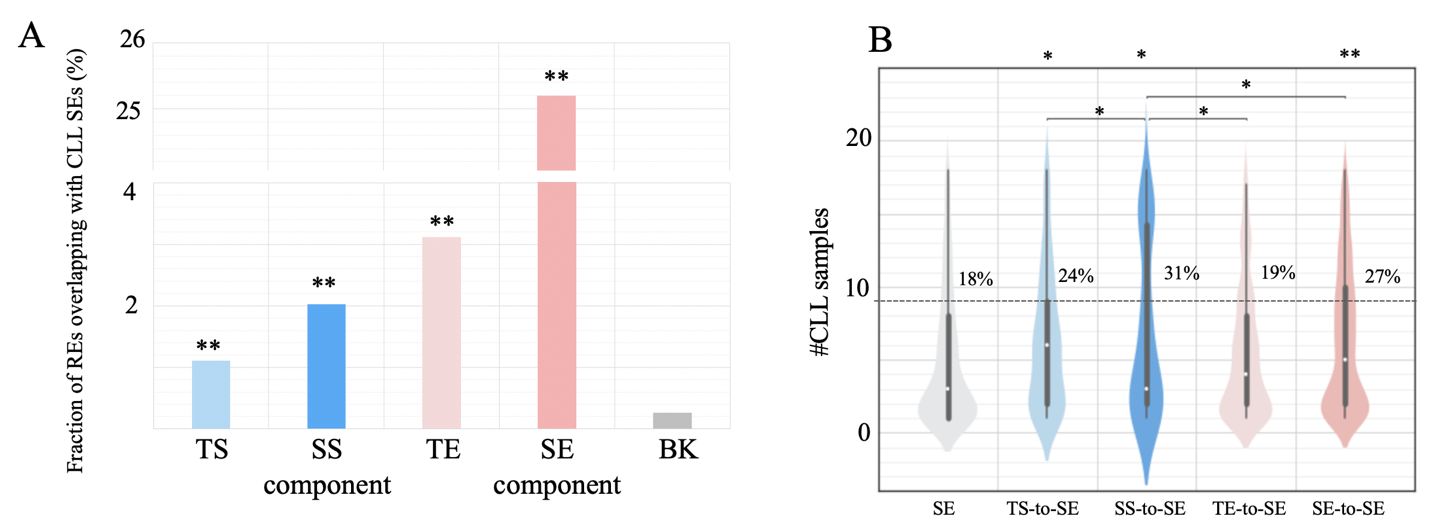

**Figure S17.** (A) Fraction of GM12878 enhancers and silencers overlapping with CLL SEs. (B) Fraction of these overlapped CLL SEs recurring across CLL samples. “BK” is DNase-seq peaks randomly selected in other cell types.

#### Supplementary Figure 18

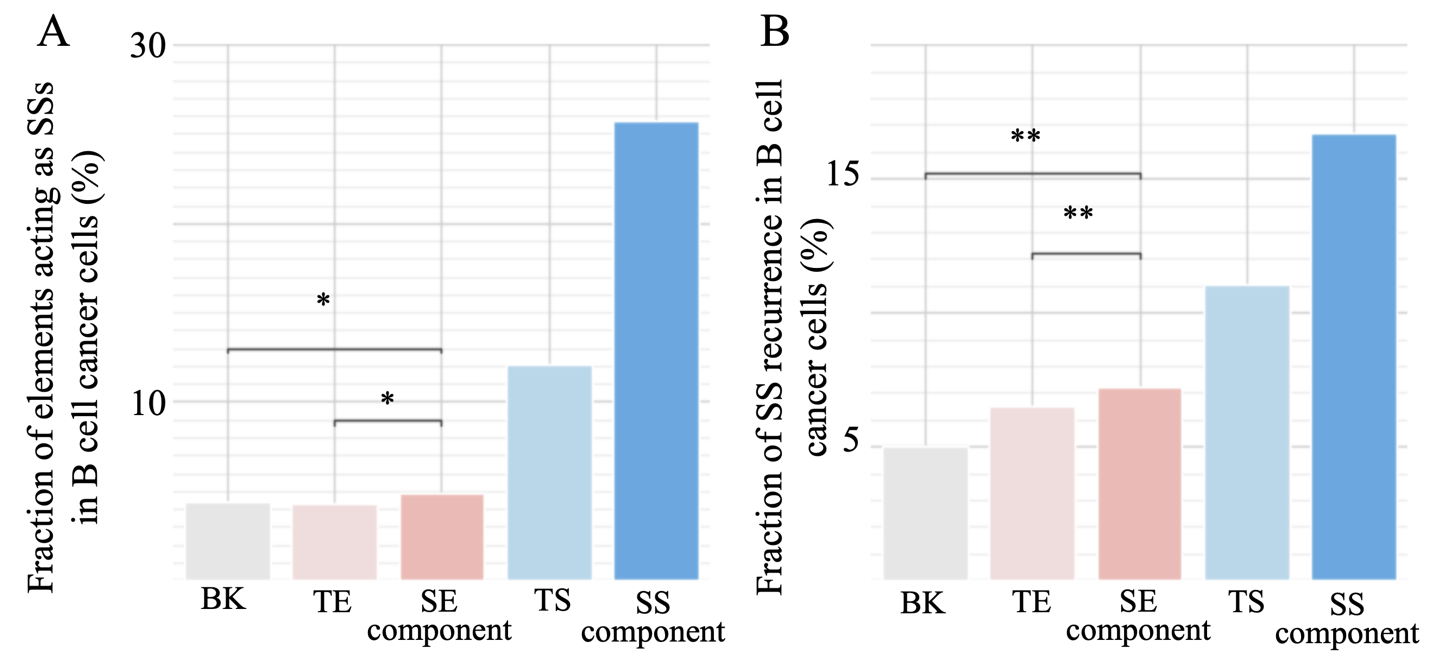

**Figure S18.** Conversions of SEs to SSs from GM12878 to B-cell cancer cells, an extreme opposite to SS-to-SEs. (A) Fraction of GM12878 enhancers and silencers overlapping with SSs in B-cell cancer cells. (B) Rate of SE-to-SSs recurring across different B-cell cancer cell lines.

$**:p<{10}^{-10}$ and $*:p\leq0.01$.

#### Supplementary Figure 19

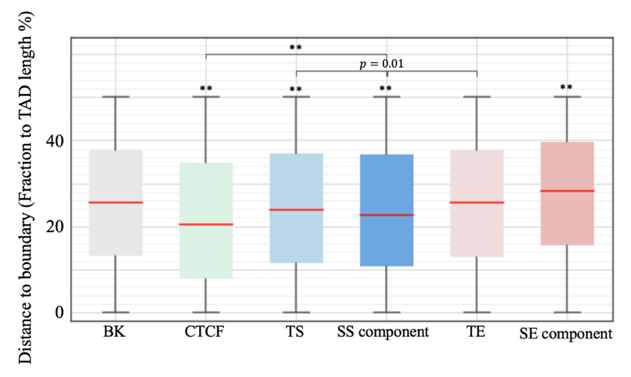

**Figure S19.** Distances of enhancers and silencers to the nearest TAD boundaries. For a fair comparison across TADs in different length, the distances to TAD boundaries are measured as fractions of the full lengths of host TADs. The asterisks above boxes indicate the significance levels in comparison to BK, i.e., DNase-seq peaks in other cell types. $**p<{10}^{-5}$.

#### Supplementary Figure 20

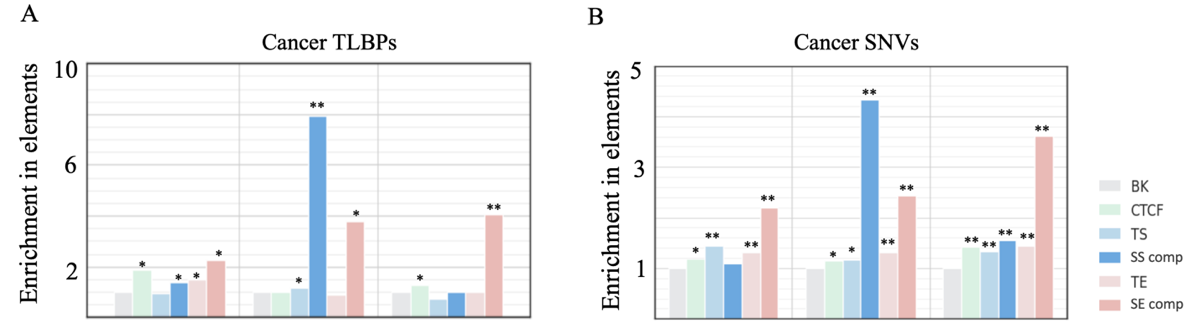

**Figure S20.** Enrichment of cancer variants in enhancers and silencers. (A) B-cell-cancer TLBPs and (B) B-cell-cancer SNVs. The enrichment levels were evaluated based on the fraction of B-cell-cancer variants among all cancer variants, to correct the biased distribution of cancer variants. “SS comp” and “SE comp” here represent SS and SE components, respectively.

$**p<{10}^{-10}$; $*p\leq0.01$.

#### Supplementary Figure 21

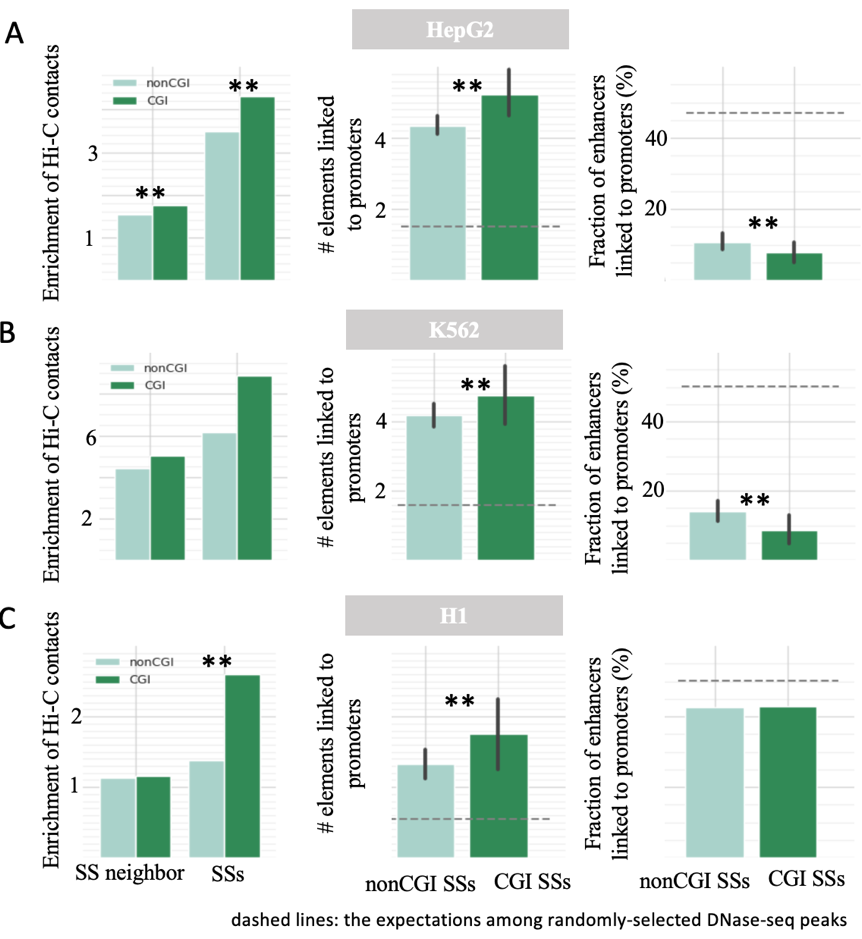

**Figure 21**. Repression models of nonCGI and CGI SSs in three cell types (A) HepG2, (B) K562, and (C) hESC H1. The dashed lines are the expected values across the whole genome. $**p<{10}^{-5}$.

#### Supplementary Figure 22

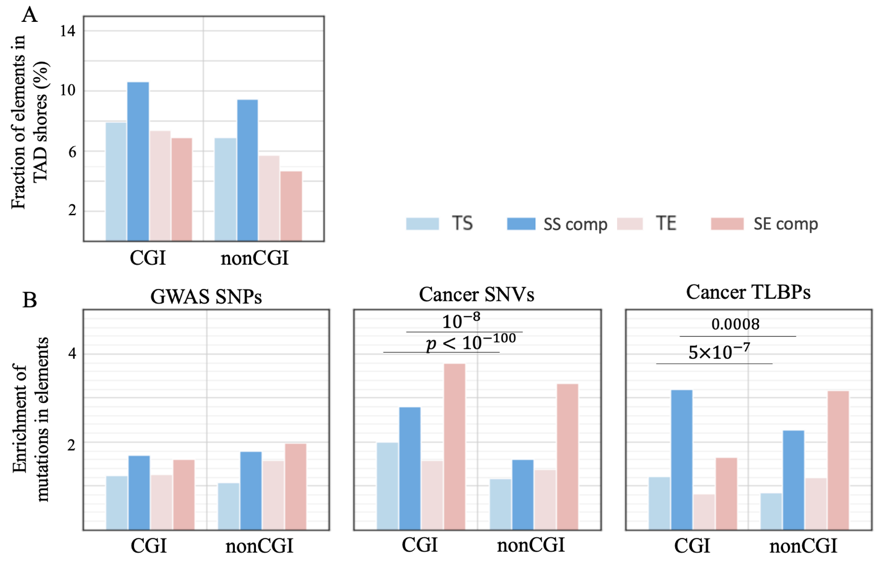

**Figure S22.** Comparisons between CGI and non-CGI SSs. (A) Fraction of elements in TAD shores. Both CGI and non-CGI SSs are enriched in TAD shores. (B) Enrichment of immunity- or B-cell-cancer- associated GWAS SNPs, B-cell-cancer SNVs, and B-cell-cancer TLBPs in CGI and non-CGI SSs. Enrichment of cancer somatic mutations were evaluated based on the fractions of B-cell-cancer mutations among all cancer mutations. “SS comp” and “SE comp” here represent SS and SE components, respectively. All these enrichment levels were evaluated as compared to DNase-seq peaks in other cell/tissue types.

**Table S1.** GM12878 SS components and TSs.

**Table S2.** SS components with top densities of B-cell-cancer SNVs.

**Table S3.** Results of luciferase experiments.

**Table S4.** Cancer samples investigated in this study.

**Table S5.** SSs and TSs in primary B cell.

**Reference：**

1	Huang, D. & Ovcharenko, I. Enhancer-silencer transitions in the human genome. *Genome research* **32**, 437-448 (2022). <https://doi.org/10.1101/gr.275992.121>

2	Whyte, Warren A. *et al.* Master Transcription Factors and Mediator Establish Super-Enhancers at Key Cell Identity Genes. *Cell* **153**, 307-319 (2013). <https://doi.org/https://doi.org/10.1016/j.cell.2013.03.035>

3	Ovcharenko, I. *et al.* Evolution and functional classification of vertebrate gene deserts. *Genome Res* **15**, 137-145 (2005). <https://doi.org/10.1101/gr.3015505>

4	Frankish, A. *et al.* GENCODE 2021. *Nucleic Acids Research* **49**, D916-D923 (2021). <https://doi.org/10.1093/nar/gkaa1087>

5	Kryuchkova-Mostacci, N. & Robinson-Rechavi, M. A benchmark of gene expression tissue-specificity metrics. *Briefings in Bioinformatics* **18**, 205-214 (2017). <https://doi.org/10.1093/bib/bbw008>

6	Cai, Y. *et al.* H3K27me3-rich genomic regions can function as silencers to repress gene expression via chromatin interactions. *Nature Communications* **12**, 719 (2021). <https://doi.org/10.1038/s41467-021-20940-y>

7	Siepel, A. *et al.* Evolutionarily conserved elements in vertebrate, insect, worm, and yeast genomes. *Genome Research* **15**, 1034-1050 (2005). <https://doi.org/10.1101/gr.3715005>

8	McLean, C. Y. *et al.* GREAT improves functional interpretation of cis-regulatory regions. *Nature Biotechnology* **28**, 495-501 (2010). <https://doi.org/10.1038/nbt.1630>

9	Sherman, B. T. *et al.* DAVID: a web server for functional enrichment analysis and functional annotation of gene lists (2021 update). *Nucleic Acids Res* **50**, W216-221 (2022). <https://doi.org/10.1093/nar/gkac194>

10	Bal, E. *et al.* Super-enhancer hypermutation alters oncogene expression in B cell lymphoma. *Nature* **607**, 808-815 (2022). <https://doi.org/10.1038/s41586-022-04906-8>
